# A Statistical Framework to Infer the Mutation Model of Tandem Repeat Variants

**DOI:** 10.64898/2026.01.21.700917

**Authors:** Luis Fernandez-Luna, Sebastián Iturbe, Carolina Adam, Nathaniel S Pope, Diego Ortega-Del Vecchyo, Rori Rohlfs

**Affiliations:** Institute of Ecology and Evolution, University of Oregon; Department of Data Science, University of Oregon; International Laboratory for Human Genome Research, Universidad Nacional Autónoma de México

## Abstract

Tandem Repeats (TRs) have complex mutational patterns that depend on many properties of the analyzed loci. An accurate characterization of the mutation model that defines the evolution of each TR is fundamental to understand the genetic diversity patterns of each TR. Here we propose a computational method that leverages the rich information contained in the ancestral recombination graph (ARG) to estimate the mutation process that drives the evolution of one loci containing a TR variant. Our method is called TRAMA, Tandem Repeat ARG-based Mutation Analysis. TRAMA uses the genealogical history estimated at each loci, which is contained in the ARG, to estimate the parameters that define the mutation of a TR under two different mutational models: The Stepwise Mutation Model (SMM) and the Two-Phase Mutation Model (TPM). First we show that TRAMA can provide estimates of the mutation rate of a TR evolving under the SMM that are accurate or have a slight underestimation when the mutation rate is higher than 10^-5^. Then, we show that TRAMA can provide reasonable estimates of the parameters that define the TPM under certain conditions. We also show that TRAMA can do an accurate selection of the mutational model that better explains the genetic diversity patterns of a loci from two competing models: TPM and SMM. Then, we show that estimates of the mutation rate under the SMM are similar when using the true genealogical history compared to using a genealogical history estimated using an ARG inference program, SINGER. We also discuss potential extensions of our methodology to perform a more accurate characterization of the mutation model driving the evolution of TRs.

## Introduction

Tandem repeats (TRs) are repetitive DNA sequences commonly classified into short tandem repeats (STRs) and variable number tandem repeats (VNTRs) depending on the motif length of their repeated units. STRs include repeat units spanning 1 to 6 base pairs. On the other hand, VNTRs feature longer motifs of 7 base pairs or more and usually encompass motif lengths up to 20 or even 50 base pairs ^1–5^. Together, these two highly mutable classes comprise approximately 8% of the human genome. The repetitive structure of TRs predisposes them to slippage events during DNA replication, resulting in changes in the number of repeat units ^5–7^. TRs exhibit mutation rates significantly higher than those of single nucleotide variants, ranging from 10^−6^ to 10^−2^ mutations per locus per generation ^8^ depending on properties of the locus associated with motif length, sequence composition, and the constancy and length of the repeat tract. For instance, TRs with consistent repeats mutate at higher rates than TRs with inconsistent repeats ^2,7,9,10^. TRs can have relevant phenotypic consequences that cause them to be under natural selection ^11^ or have important clinical outcomes due to their contribution to different diseases such as Huntington’s disease and fragile X syndrome ^12,13^.

Short read sequencing has limitations in accurately resolving these complex regions as TRs often approach or exceed the length of the read ^12,14^. Thus, TRs have been historically overlooked due to technical challenges in genotyping, even after the advent of next-generation sequencing. However, recent advances in long-read sequencing have improved our ability to resolve TRs’ structure and distribution, providing deeper insights into their biological roles ^12,14,15^. This has led to significant advancements in understanding TRs role in the modulation of gene expression through their effect on methylation patterns ^16–18^, transcription factor binding affinity ^19,20^ and changes on RNA or protein structure ^21–23^. Our ability to genotype TRs opens the door for more careful analyses to understand their evolution and their impact on various phenotypes.

Another important recent advance in population genetics is the recent development of methods to estimate the genealogical history at each locus in the genome. The genealogical history at each locus is part of a structure defined as the ancestral recombination graph (ARG) ^24^. An ARG captures the full evolutionary history of a locus and has the complete set of ancestral relationships among haplotypes. The development of ARG estimation methods has allowed an influx of new methods that leverage the genealogical history at each locus to provide more accurate inferences of past evolutionary processes than methods that do not directly use genealogical information^25^. Here we develop TRAMA, Tandem Repeat ARG-based Mutation Analysis, to gain deeper insights into the evolutionary processes shaping TR variation. Our approach uses a novel maximum likelihood framework that leverages the genealogical history contained in the ancestral recombination graphs (ARG) to infer the mutational model acting on each TR loci. We aim to more accurately estimate mutation rates and the underlying mutation model acting at each TR loci by conditioning on the genealogy of that TR locus.

Here we show how we can use TRAMA to estimate the parameters that define two mutational models: the Stepwise Mutation Model (SMM) and the Two-Phase Mutation Model (TPM). We show that TRAMA can estimate the mutation rate of a TR loci under the SMM accurately or with a slight underestimation. Then, we show that TRAMA can accurately estimate the parameters of the TPM under certain conditions. Finally, we show that our method can be used to perform model selection between the two competing models of SMM and TPM. Broadly, we show that our TRAMA can leverage population genetic data to infer the mutation model underlying each TR locus.

## Results

### Estimates of mutational model parameters

We analyzed the accuracy of TRAMA to estimate the TR mutation rate evolving under the SMM using simulations where we know the true genealogy of each TR locus. We performed simulations of TRs with five different sample sizes. We find that the estimation of the mutation rate (μ) in TRs evolving under the Stepwise Mutation Model (SMM) is accurate or slightly underestimated in TRs with a mutation rate μ = 10^-3^, 10^-4^ and 10^-5^ under three different sample sizes (Figure 1). In contrast, for TRs with a lower mutation rate of μ = 10^-6^ we obtained inaccurate underestimates of more than 2 orders of magnitude.

**Figure 1.**
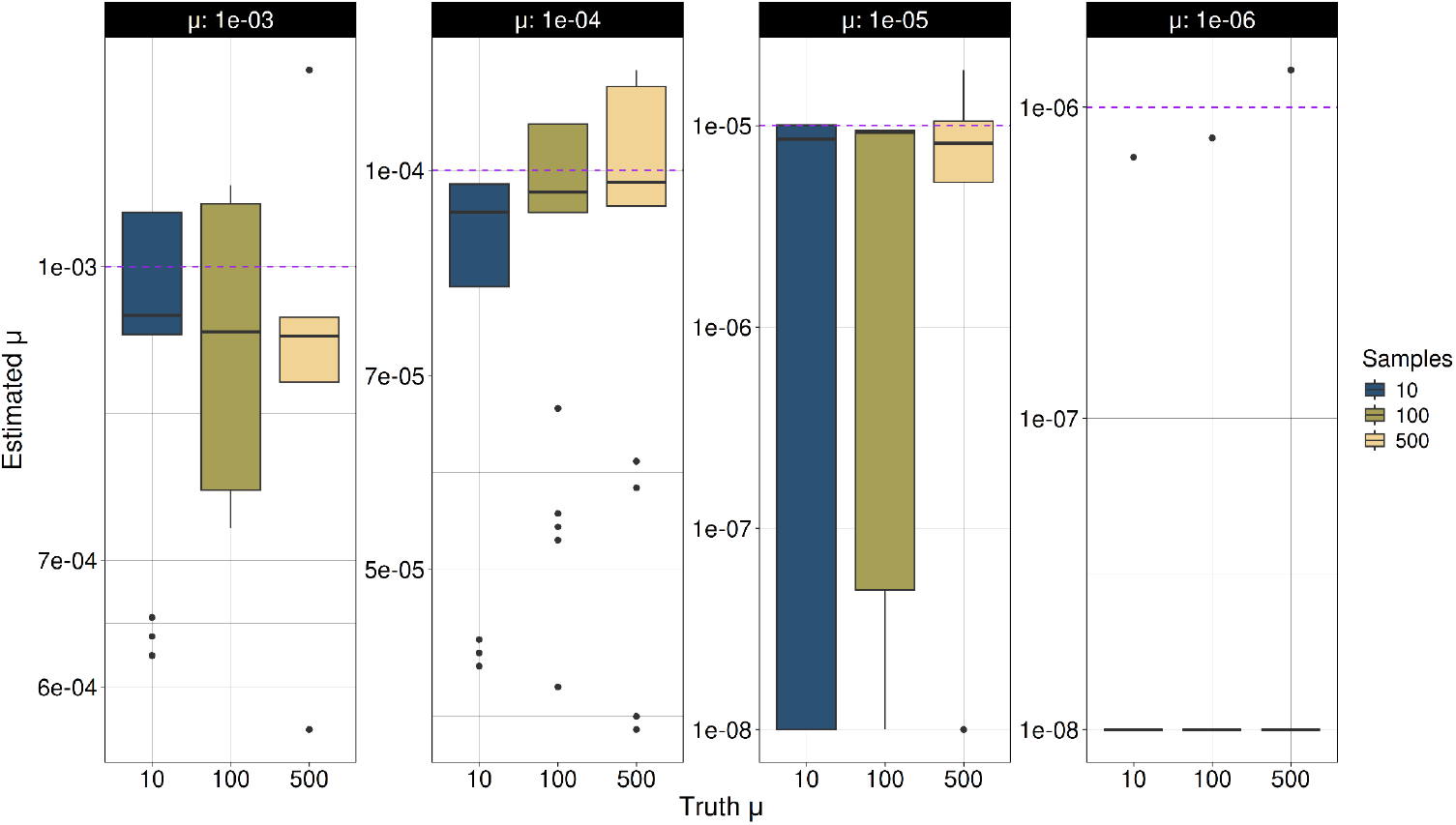
Estimation of μ under the SMM. Tandem repeat (TR) variants were simulated in independent simulations for each mutation rate (μ = 10^−3^ to 10^−6^) under the SMM. For each simulated dataset, μ was estimated using TRAMA across three different sample sizes (N = 10, 100 and 500 individuals).

We also analyzed the accuracy of the mutation process parameters estimated with TRAMA under the Two-Phase Mutation Model (TPM) (Figure 2A, S1). We find that estimates of the mutation rate (μ) are accurate or present a slight underestimation across multiple parameter values of *p* and *m* at using different sample sizes (Figure 2A; Figure S1A). Estimates of parameters *p* and *m* are more sensitive to μ, with high accuracy for high mutation rate, but lower accuracy for lower mutation rates (Figure 2B-C; Figure S1B-S1C). Estimates of *p* and *m* are additionally sensitive to the underlying values of each other, with decreased accuracy for higher values of *m* and *p*, respectively.

**Figure 2A.**
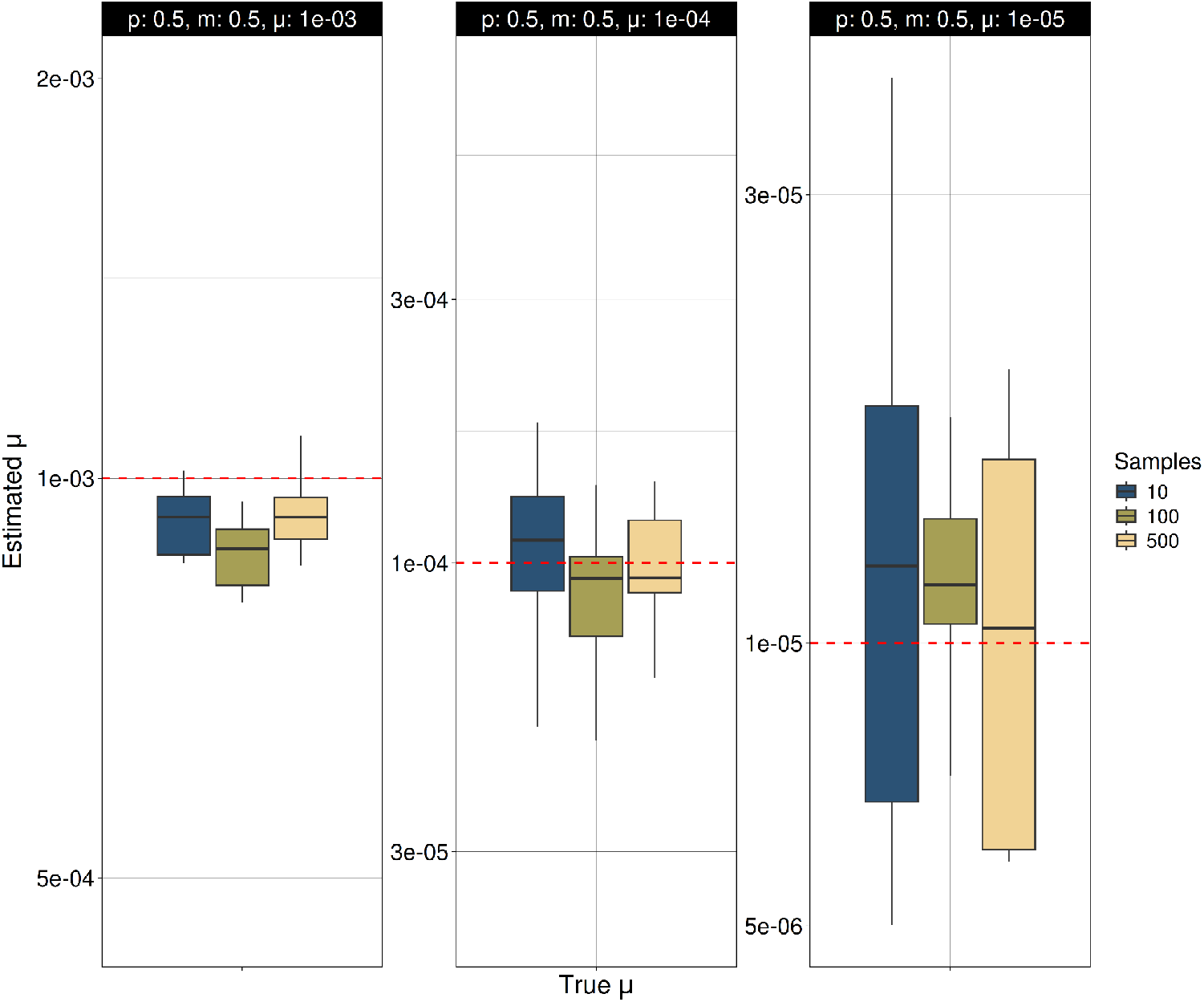
Estimation of the mutation rate (μ) under the Two-Phase Mutation Model (TPM). Tandem repeat (TR) variants were simulated under the TPM in independent simulations across different mutation rates (μ = 10^−3^ to 10^−5^). Mutation rate (μ) was estimated using TRAMA across multiple sample sizes (N = 10, 50, 100 and 500), conditioning on the true ancestral recombination graph (ARG). The TPM parameters were held constant at *p* = 0.5 and *m* = 0.5.

**Figure 2B.**
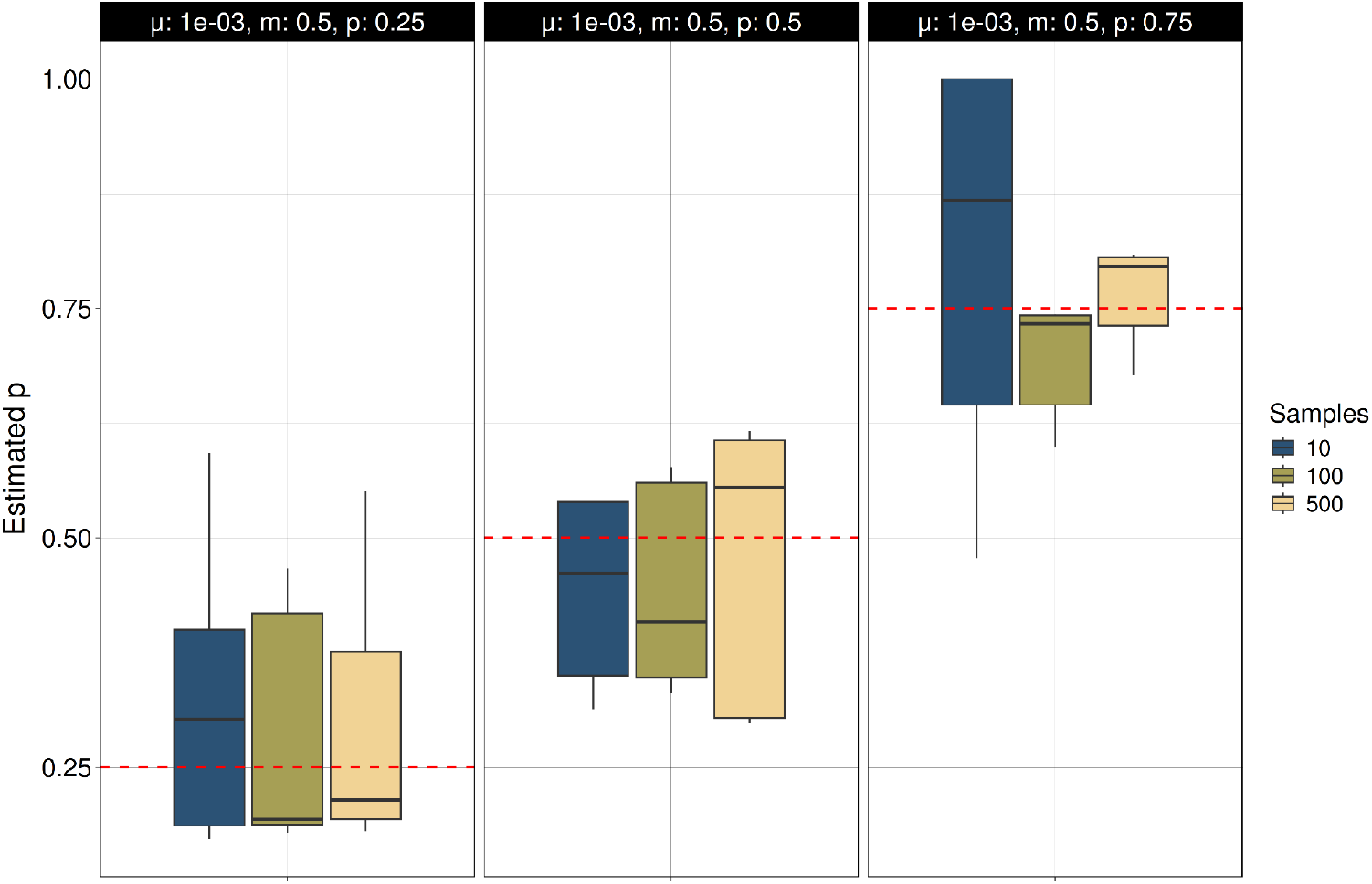
Estimation of the probability of multistep mutations (*p*) under the Two-Phase Mutation Model (TPM) using the true ARG. TR variants were simulated under the TPM in independent simulations across different mutation rates (μ = 10^−3^ to 10^−5^). The probability of multistep mutations (*p*) was estimated using TRAMA across multiple sample sizes (N = 10, 100 and 500), conditioning on the true ARG. Simulations were performed with fixed values of μ= 0.001 and *m* = 0.5.

**Figure 2C.**
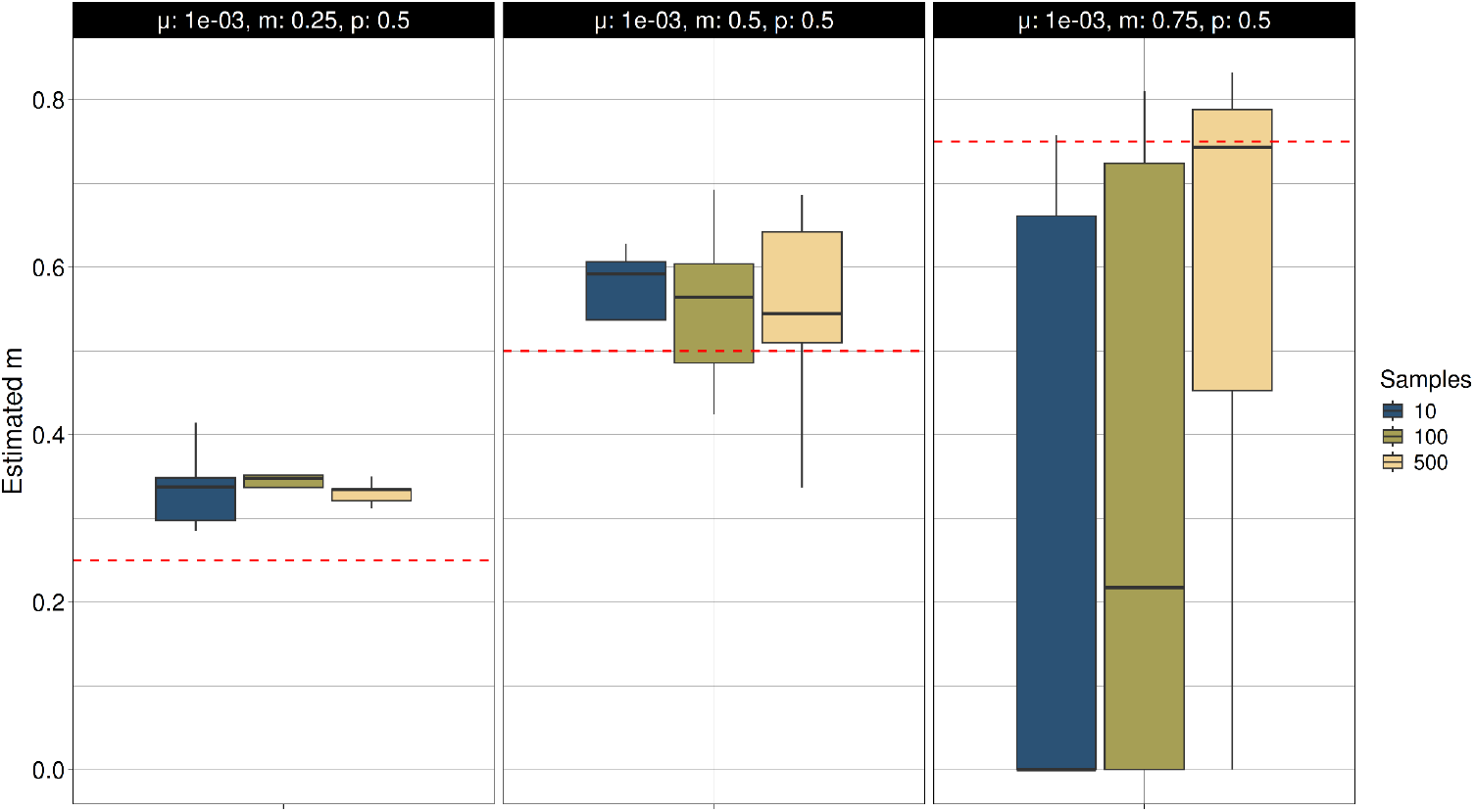
Estimation of the multistep mutation length parameter (*m*) under the Two-Phase Mutation Model (TPM) using the true ARG. TR variants were simulated under the TPM in independent simulations across mutation rates (μ = 10^−3^ to 10^−5^). The multistep mutation length parameter (*m*) was estimated using TRAMA across multiple sample sizes (N = 10, 100 and 500), conditioning on the true ARG. Simulations were performed with fixed values of *p* = 0.5 and μ=0.001.

### Model comparison

We analyzed how TRAMA can classify the mutational model that better explains the evolution of a single TR loci between two competing models: SMM and TPM (see methods: Assess of mutational model classification). With a high mutation rate (μ=10^-3^), TRAMA correctly classifies TRs evolving under the SMM in more than 97.5% of the simulations with sample sizes of 10, 100 and 1,000 individuals (Figure 3). The correct classification of TRs evolving under the SMM improves slightly with larger sample sizes (0.983, 0.990, and 0.987 for n = 10, 100, 1000, respectively). With a lower mutation rate (μ=10^-5^), TRAMA correctly classifies 72.8%, 98.4%, and 97.8% of the TRs evolving under the SMM with sample sizes of n=10, 100, and 1000, respectively. (Figure S2A).

**Figure 3.**
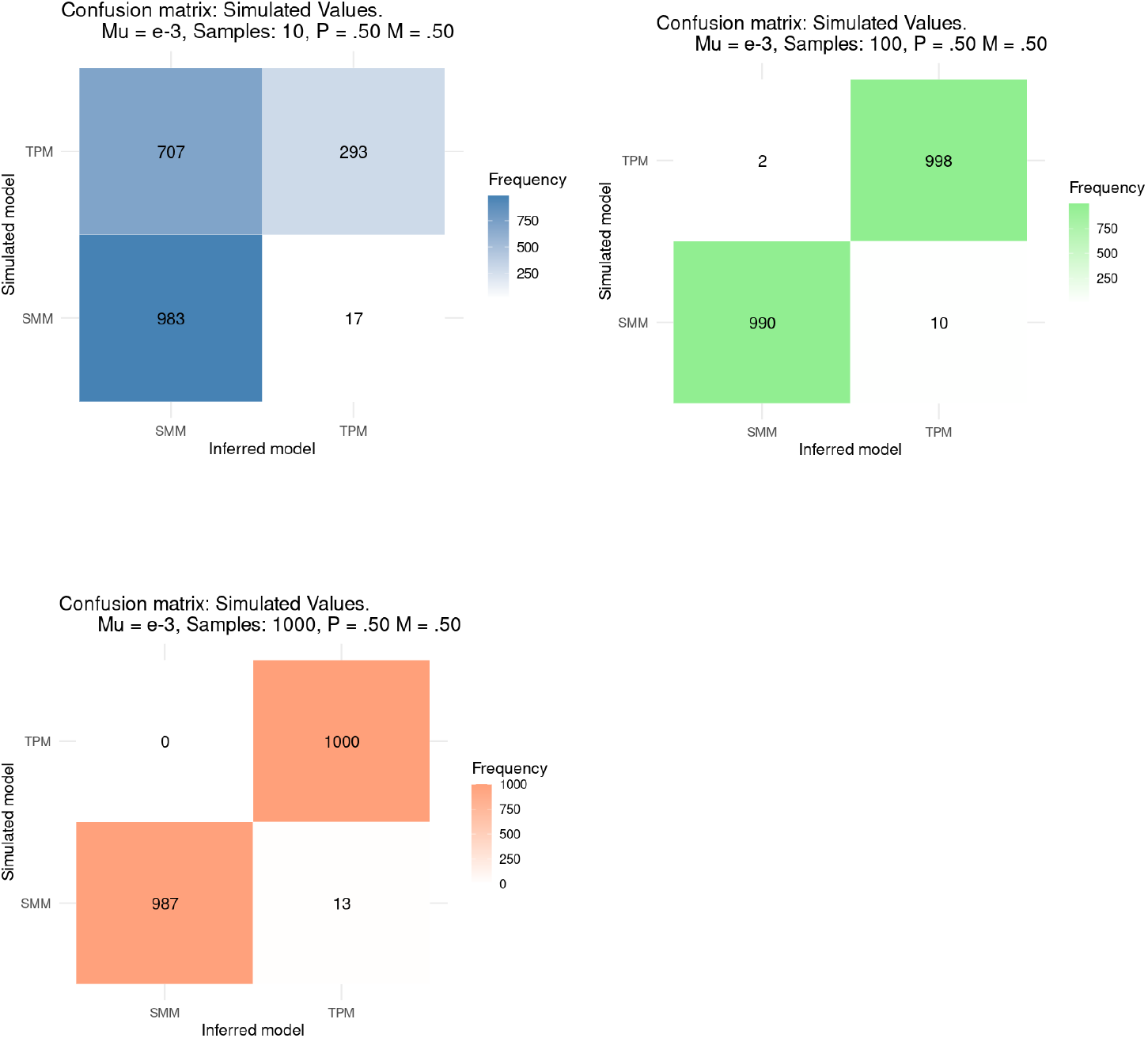
Confusion matrices for 3 different sample sizes (N = 10, 100 and 1000) showing how TRAMA classifies 1,000 simulations as evolving either under the SMM or TPM.

We also analyzed the ability of TRAMA to classify the mutational model of TRs evolving under the TPM with parameters p = m = 0.5. In this case, with a high mutation rate (μ=10^-3^), sample sizes have an important effect. TRAMA can accurately classify a TR as evolving under the TPM in only 29.3% of the simulations when the sample size is equal to 10. TRAMA can accurately classify a TR as evolving under the TPM on 99.8% and 100% of simulations when the sample size is equal to 100 and 1000, respectively. TPM inference accuracy remains high with frequent multistep mutations (low *p*) and longer multistep mutations (low *m*). However accuracy is lower with less frequent multistep mutations (high *p*) and smaller multistep mutations (high *m*) (Figure S2B). Accuracy is also lower with lower mutation rates, as well as smaller sample sizes (Figure S2C).

### Impact of using inferred genealogy

The results presented in the past sections of the paper condition on the correct genealogy at each TR locus analyzed. However, the genealogy at each TR loci must be inferred and inaccuracy of that inference and could cause errors in mutational model parameter estimates. We analyzed how the use of a genealogy inferred using SINGER ^26^ an ARG inference method, impacts the accuracy of the estimated mutation model parameters. Although inaccuracies in ARG inference reduce estimation precision, their overall impact is modest (Figures 4, S3). Mutation rate, μ, under the SMM is consistently estimated with more accuracy and precision when conditioning on the true genealogy rather than the inferred genealogy in simulations (Figure 4). Broadly, estimates of μ are accurate or have a slight overestimation when using the inferred genealogy when μ is bigger than 10^-5^ (Figure S3). The estimates of μ also improve with a larger sample size.

**Figure 4.**
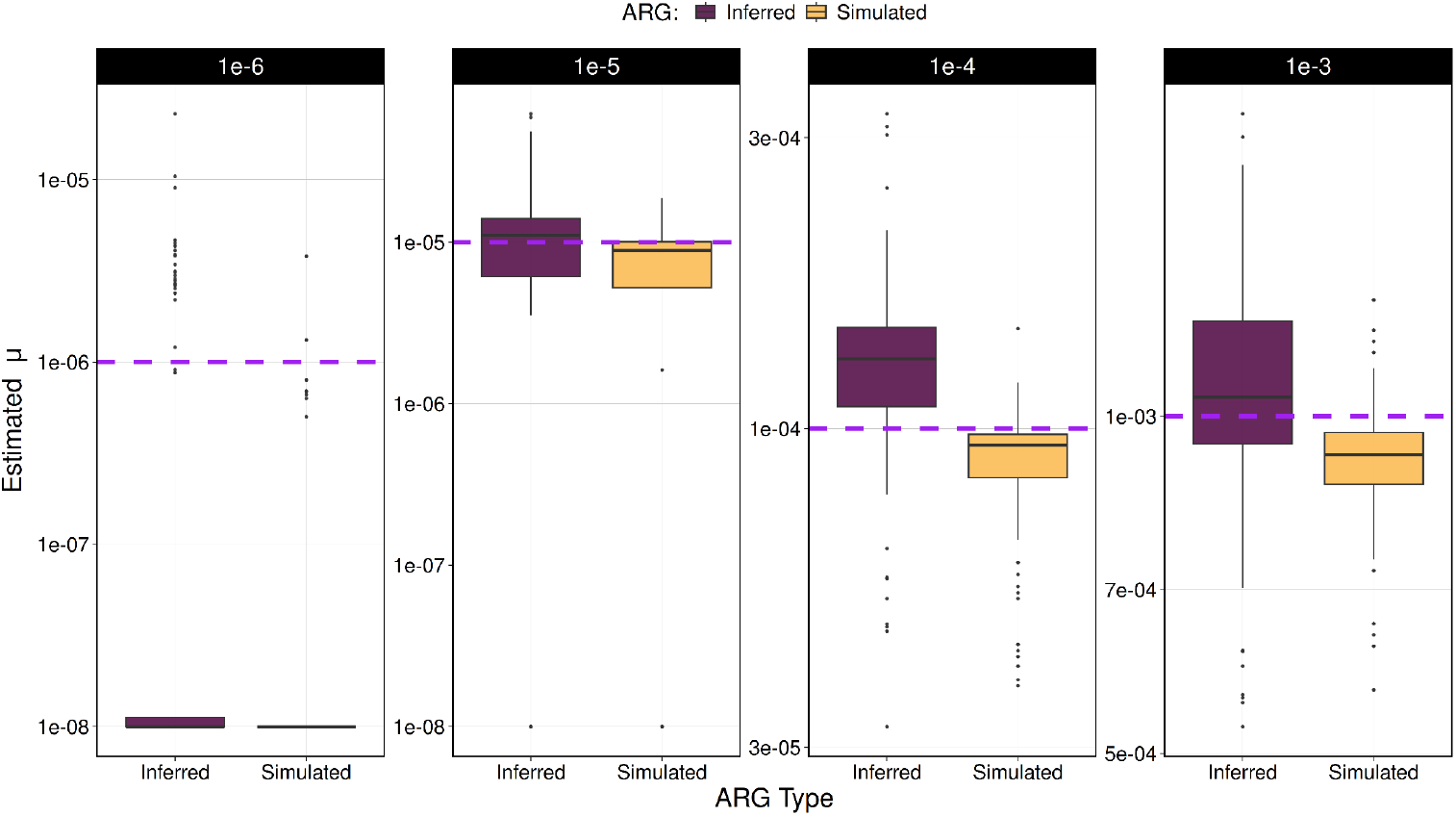
Comparison of mutation rate (μ) estimation under the Stepwise Mutation Model (SMM) using true versus estimated genealogies. Tandem repeat (TR) variants were simulated in independent simulations for each mutation rate (μ = 10^−2^ to 10^−6^) under the SMM. Mutation rate (μ) was estimated using TRAMA conditioning on either the true genealogy or a genealogy inferred from the data. Estimates conditioned on the estimated genealogy (inferred) exhibit increased variance relative to those based on the true genealogy (simulated).

Estimates of μ, *p*, and *m* under inferred genealogies exhibit greater variability than those that use the true genealogy (Figure S4A-C). We also note that conditioning on the true genealogy tends to provide somewhat more accurate and precise estimates of μ, *p*, and *m* (Figure S3). However, we see that estimates of μ, *p*, and *m* that use the inferred genealogy tend to have a larger variance but do not appear to be more biased than those that use the true genealogy (Figure S3). We particularly see that estimation of the mutation rate, μ, under the TPM remains stable and accurate across variations in *p, m*, and sample size, even when using the inferred ARG (Fig. S4A). Estimates of *p* and *m* improve when the sample size increases under the TPM with certain parameter values of *p* and *m* using the estimated genealogy (Figure S4B-C).

## Discussion

Recent studies have found that some methods that leverage the ARG provide more accurate inferences of past evolutionary processes than methods that do not leverage the ARG ^24,25,27^. These accurate inferences in methods that leverage the ARG are due to the rich evolutionary information contained in the ARG. Here we developed a new approach, TRAMA, to infer the TR mutational model by leveraging the information contained in the ARG. TRAMA infers the mutational model with reasonable accuracy when mutation rates are high, creating more mutational data to power the inference. The inference of the mutational model becomes less accurate with lower mutation rates, and especially in a TPM where multi-step mutations are infrequent (high *p*) and where multi-step mutations are small (high *m*). With those parameter values, the TPM is more SMM-like, making the models difficult to distinguish. We find that TRAMA can provide estimates of the mutation rate that are on the same order of magnitude of the actual value or that are slight underestimations. Additionally, we find that TRAMA can provide accurate estimates of the mutation rate μ and the parameters *p* and *m* of the two phase mutation model when with similar parameters that provide higher accuracy in model inference. Additionally, we find that inferences and estimates improve with larger sample sizes.

TRAMA uses information from the sequence of genealogies across the genome to infer the parameters of the mutation rate model. However, the sequence of genealogies needs to be inferred from the SNP data, and its inference can have inaccuracies that can cause problems with the estimation of parameters from the TR mutation model. Our results show that estimates of the mutation rate are concordant when using the true sequence of genealogies and the sequence of genealogies inferred from SNP data using SINGER. These results are encouraging since they tell us inaccuracies in ARG inference do not dramatically impact inference of TR mutational model or parameters.

Overall, we present a new method, TRAMA, that is capable of estimating parameters of the TR mutation model and estimating the TR mutation model that better explains the patterns of genetic variation seen in the data. TRAMA presents a framework that can be extended to analyze other TR mutational models and to analyze the complex mutational dynamics of TRs at a genomic level using population genomic data.

## Methods

### Inferring mutational model and estimating parameters with TRAMA

We developed TRAMA, a statistical tool to estimate mutational model parameters of individual TRs, and to determine their individual mutational model: SMM or TPM (Figure 5). TRAMA uses a maximum likelihood framework to estimate the mutational model parameters of observed TR alleles conditioning on the genealogical relationships at the observed locus. Under the SMM, TRAMA estimates the mutation rate, μ,. Under the TPM, TRAMA estimates μ, the probability of a stepwise mutation (*p*), and the parameter of the geometric distribution describing the size of multistep mutations (*m*).

**Figure 5.**
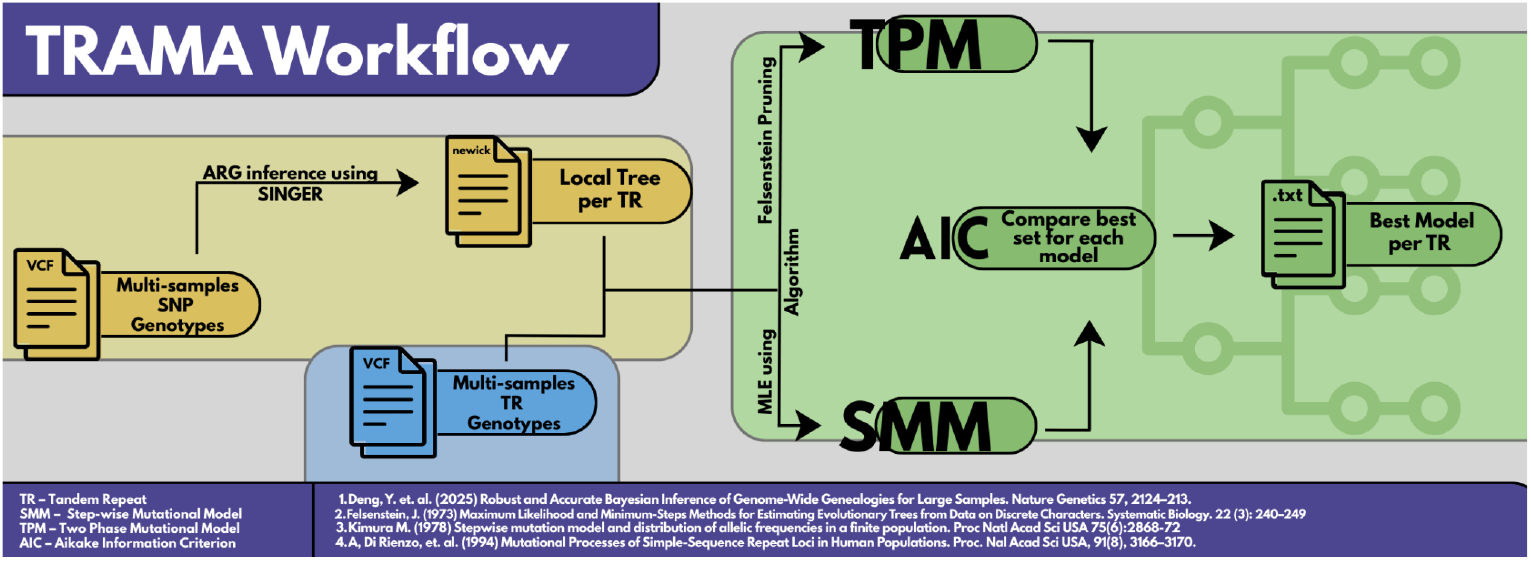
TRAMA workflow. This figure illustrates the complete TRAMA pipeline for inferring and comparing tandem repeat (TR) mutation models. Multi-sample TR genotypes and SNP genotypes, provided as VCF files, are used as inputs. SNP genotypes are first analyzed with SINGER to infer an ancestral recombination graph (ARG), from which a local genealogical tree (in Newick format) is extracted for each TR locus.

### Maximum likelihood estimation of TR mutation model parameters using genealogies

We used the Felsenstein pruning algorithm (FPA) to estimate the maximum likelihood parameter values for a particular mutational model. The FPA is a dynamic programming algorithm to compute the likelihood of a set of observed data on a phylogeny under a specified model of evolution^28^.

Conditioning on the trees, we model the evolution of short tandem repeat (STR) genotypes observed for each sampled individual. STR genotypes are treated as observed character states evolving along the branches of the local tree.

The FPA assumes that each observable character across samples belongs to a finite set of discrete states. In TRAMA, we adopt the same assumption but redefine the state space such that each state corresponds to a motif repeat length of an STR allele. Transitions between repeat-length states along each branch are governed by a continuous-time Markov chain (CTMC).

Under this framework, the evolution of STR states along a branch of length (t) is described by a transition probability matrix

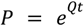

where Q is the transition rate matrix specified by the chosen STR mutation model.

Likelihood computation proceeds by recursively evaluating partial likelihoods at internal nodes of the tree using the FPA. For an internal node v with two descendant nodes (c1) and (c2), the partial likelihood of observing the STR genotype data in node (v) being in repeat-length state (i), is given by

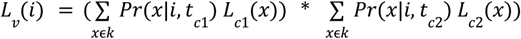

Here, Lv(i) denotes the likelihood of having the STR genotype i in the node v. The sums are taken over all possible STR repeat-length states (x) in the defined space. The term Pr(x∣i,tck) represents the transition probability that an STR allele in state i at node v mutates to state x at descendant node ck over a branch of length *tck*. These transition probabilities are obtained from the continuous-time Markov chain describing STR evolution and are computed from the transition probability matrix P(t)=exp(Qt), where Q is the model-specific transition rate matrix.

The quantities Lc1(x) and Lc2(x) are the partial likelihoods at the descendant nodes, conditional on state x, and summarize the likelihood of the observed STR genotypes in the subtrees rooted at c1 and c2, respectively.

This recursive formulation integrates over all possible ancestral STR states at internal nodes, thereby marginalizing over unobserved repeat-length configurations. At the tips of the tree, partial likelihoods are initialized such that Lv(i)=1 if the observed STR genotype corresponds to state i, and Lv(i)=0 otherwise.

At the root of the tree, the likelihood of the observed STR genotypes is obtained by summing over all possible root states. This procedure does not require explicit reconstruction of ancestral repeat lengths.

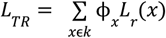

Under this formulation, inference of the maximum likelihood estimator depends exclusively on two components: the inferred local tree (topology and branch lengths) obtained from the ARG, and the STR mutation model, which determines the structure of the transition rate matrix Q. All information about mutational dynamics is therefore encoded in the model parameters, while the genealogical history is treated as fixed and known. We find the parameter values that maximize *L*_*TR*_ using the optimization algorithm L-BFGS-B.

### STR mutation model and the transition rate matrix

TRAMA currently supports two mutation models for short tandem repeat (STR) evolution: the stepwise mutation model (SMM) and the two-phase mutation model (TPM).

The SMM, originally proposed by Kimura and Ohta^29^, is one of the most widely used models for microsatellite mutation. Under the SMM, the STR motif repeat length changes by exactly one unit when a mutation occurs, either increasing or decreasing with equal probability.

To account for more complex mutational behavior, TRAMA also implements the TPM, introduced by Di Rienzo et al.^30^. The TPM assumes that STR mutations occur either as single-step changes or as multistep changes. A mutation results in a single-step change in repeat length with probability p. The mutation occurs as a multistep event with probability 1−p. A multistep event allows the repeat length to change by more than one unit in a single generation. Mutations occur at a constant rate μ independent of the current repeat length under both models.

The SMM and the TPM mutation models are incorporated into TRAMA by defining an appropriate transition rate matrix (Q), which specifies the instantaneous rates at which a continuous-time Markov chain (CTMC) transitions between repeat-length states.

### Transition rate matrix under the SMM

The SMM yields the simplest form of the transition rate matrix. The matrix depends only on the mutation rate μ and the number of allowed repeat-length states N. Because the SMM restricts mutations to single-step changes, each state can transition only to its instant neighboring states i−1 and i+1.

Under the SMM, the transition rate matrix is defined as

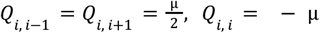

with all other entries equal to zero. This construction ensures that each row of the matrix sums to zero, as required for a valid CTMC (Table 1).

**Table 1.**
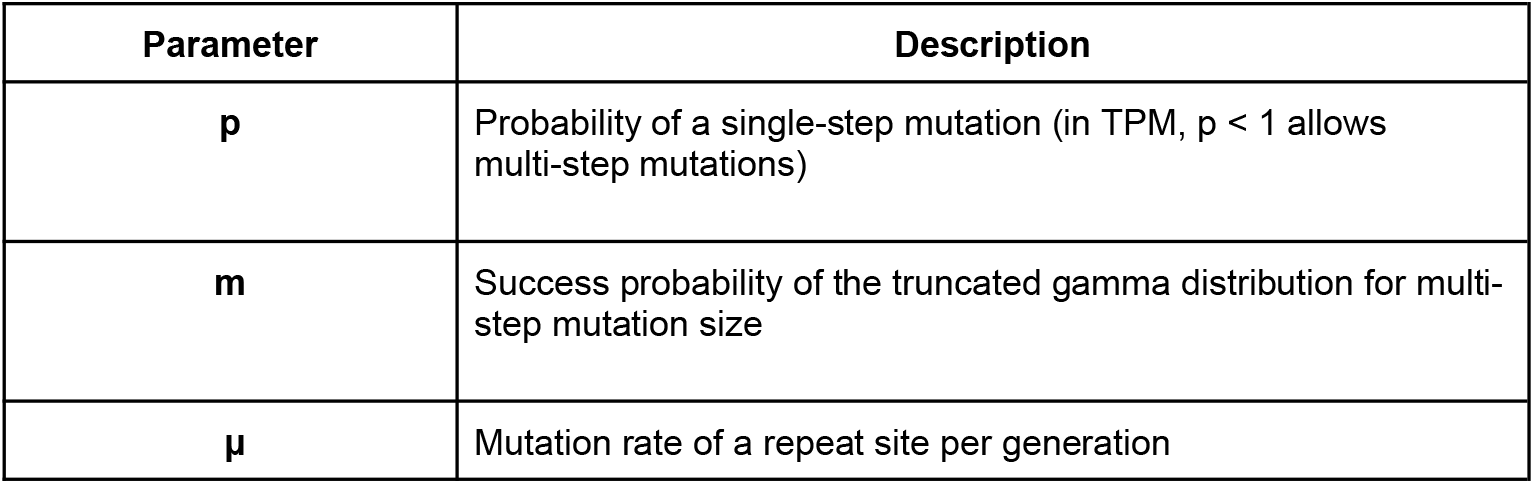
Description of parameters used in Sainduiin^31^.

### Transition rate matrix under the TPM

The TPM introduces additional flexibility by allowing both single-step and multistep mutations. In this model, the parameter p specifies the probability that a mutation event results in a single-step change. When a multistep mutation occurs, the size of the repeat-length change is governed by a truncated gamma distribution.

To implement the TPM within a finite state space, the continuous gamma distribution must be discretized and truncated to the set of allowable repeat-length changes. Following Sainudiin et al.^31^, the probability of a multistep mutation resulting in a change of k repeat units is denoted by γ(m), where m controls the shape of the geometric distribution of the multistep process.

Under the TPM, the off-diagonal entries of the transition rate matrix are given by:

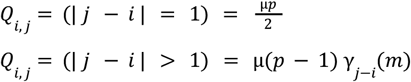

with diagonal entries defined such that each row sums to zero. Here, γ∣j−i∣(m) represents the probability mass assigned to a multistep change of size ∣j−i∣under the given distribution.

Under this formulation, the TPM transition rate matrix depends on three parameters: the mutation rate μ, the probability of single-step mutations p and the parameter m which controls the relative frequency of larger mutational jumps.

### Transition Probability Matrix

The transition probability matrix is directly derived from the transition rate matrix and defines the probabilities of transitioning between repeat-length states over a given time interval (t). This matrix is square with dimension N×N, where N denotes the number of discrete STR repeat-length states. Each entry Pij(t) represents the probability that the system transitions from state i to state j over evolutionary time t. In the Felsenstein pruning algorithm, the transition probability matrix quantifies the probability of mutating from any given repeat-length state to any other state, conditional on the divergence time between nodes.

### Simulation Framework to test the parameter estimates of TRAMA

To assess the accuracy of TRAMA, we applied it to simulated tandem repeat (TR) and SNP haplotypes. Simulations were performed using SLiM v4.1^32^ under a single-population model with a constant effective population size (Ne = 10,000) and a total sequence length of 10 Mb. A total of 1,000 TR loci were simulated and randomly distributed along the genome.

We modeled TRs with mutation rates distinct from those of SNPs, reflecting the known heterogeneity in TR mutation rates arising from differences in sequence composition, motif length, and allele size. The per-base, per-generation SNP mutation rate (μ_snp_) and recombination rate (r) were both set to 1.2 × 10^−8^. Repeat expansions and contractions were simulated using the mathematical framework developed by Sainudiin^31^, under two evolutionary models: the Stepwise Mutation Model (SMM) and the Two-Phase Mutation Model (TPM).

Under the SMM, TR mutation rates (μ_tr_) were set to 10^−3^, 10^−4^, 10^−5^, and 10^−6^. Under the TPM, μ□_r_ was varied similarly, and we additionally explored values of the probability of multistep mutations (*p* = 0, 0.25, 0.5, 0.75) and the multistep mutation length parameter (*m* = 0, 0.25, 0.5, 0.75). Simulations were conducted across combinations of these parameters to evaluate their joint effects on TRAMA’s accuracy.

Finally, we assessed the impact of sample size on parameter estimation by varying the number of sampled individuals (N = 10, 50, 100, 150, 200, and 1,000).

### Assessment of mutational model classification

We evaluated the performance of TRAMA for discriminating between the two tandem repeat mutational models: the SMM and TPM. In order to do this, we used TRAMA to analyze TRs which were generated using msprime 1.0^33^ under a constant population size of 10,000 individuals and a total sequence length of 1 Mb. We set the SNP mutation rate and the recombination rate to be 1.2 x 10^−8^ per base per generation. We use the Akaike Information Criterion (AIC) to compare mutational model for each locus:

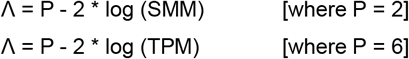

### Genealogy estimates

TRAMA first uses SINGER^26^ to infer the genealogy of a TR centered on the TR in the center of a region of 10 Mb. TRAMA uses the SNP data from mutations surrounding a TR to estimate the sequence of genealogies across the 10 Mb window. Then, we impute the genealogy at the TR loci to estimate using the genealogical data surrounding the TR loci. SINGER was implemented using the following parameter: SNP mutation and recombination rates were set to μ = r = 1.2×10−8 per base per generation, with an effective population size Ne = 10,000. We retained 100 ARG samples by thinning every 20th iteration of the MCMC chain.

## Acknowledgements

We thank Peter L. Ralph for useful discussions on ARG inference methods. RR, LGFL and SIP were supported by the NSF CAREER 2325466 grant. DODV and SIP were supported by the NIH grant R01HG012605.

