## Supplementary Material for "A Statistical Framework to Infer the Mutation Model of Tandem Repeat Variants"

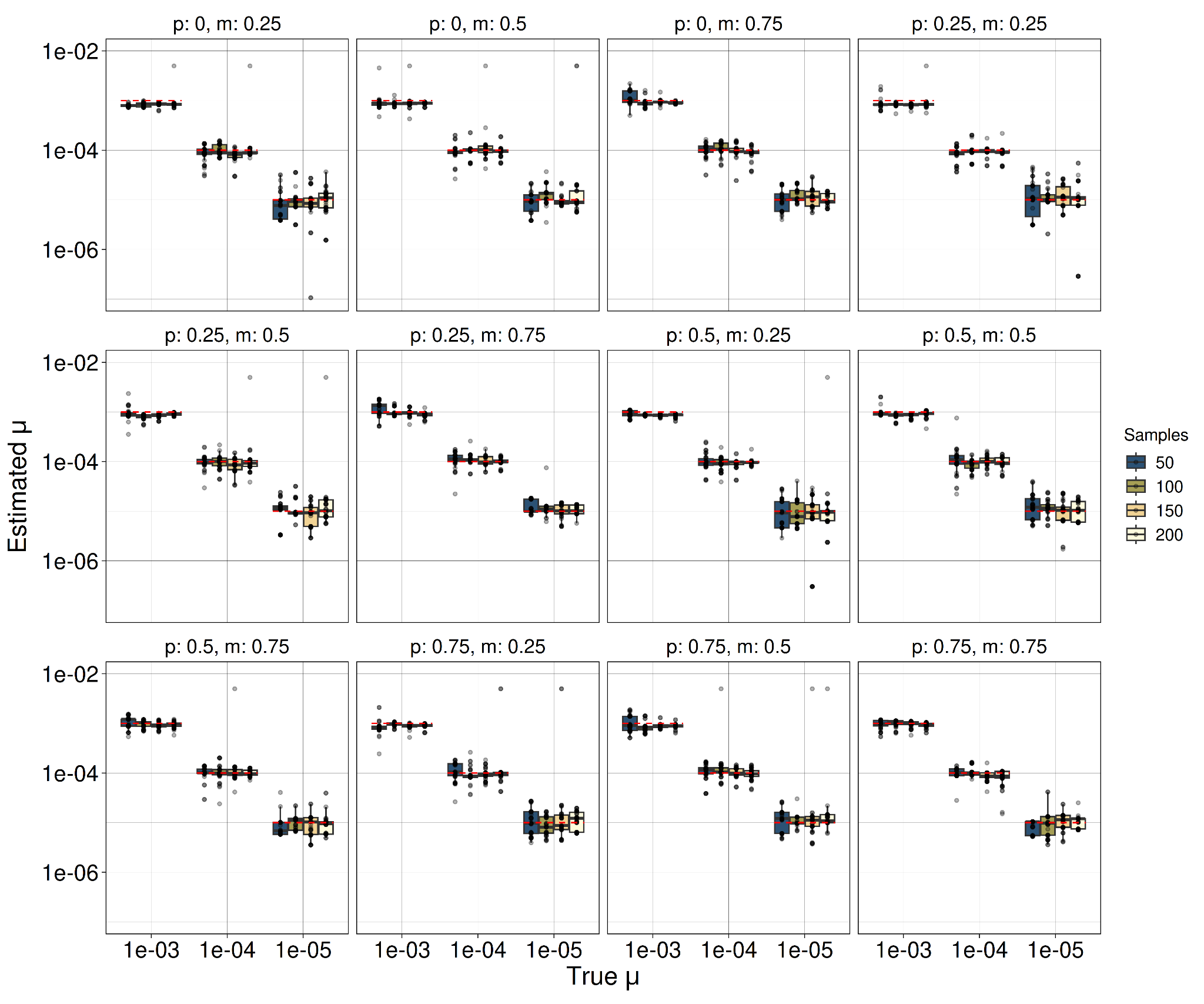

**Figure S1A. Intersectional accuracy of mutation rate (μ) estimation under the Two-Phase Mutation Model (TPM).** Mutation rate (μ) estimation accuracy was evaluated across different values of μ, *p*, *m*, and sample sizes under the TPM. TR variants were simulated in independent simulations, and μ was estimated using TRAMA while conditioning on the true ARG.

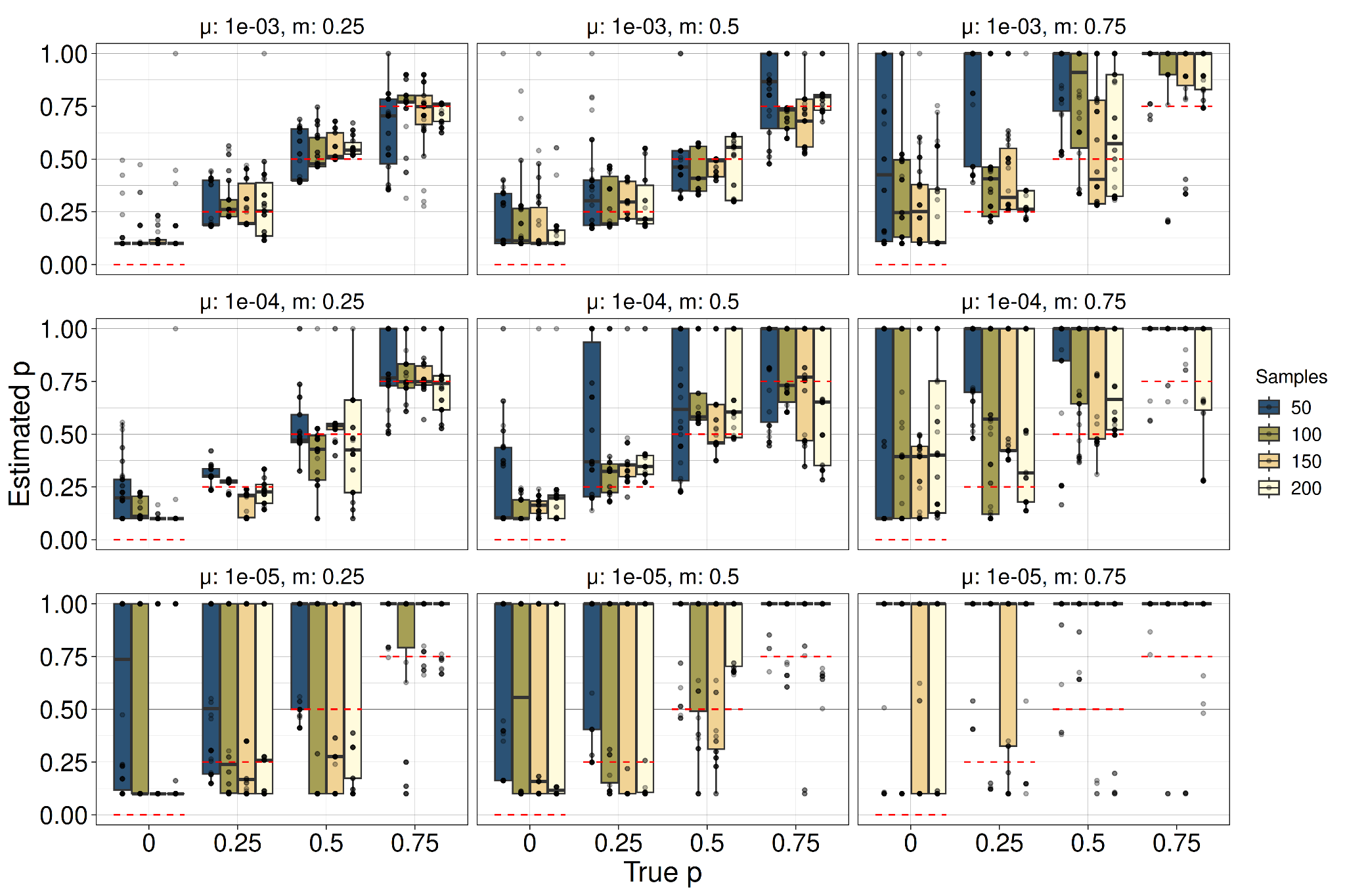

**Figure S1B. Intersectional accuracy of the probability of multistep mutations (*p*) under the Two-Phase Mutation Model (TPM).** Accuracy of *p* estimation was assessed across combinations of values of the mutation rate (μ), multistep mutation probability (*p*), multistep mutation length (*m*), and sample sizes. Parameter estimates were obtained using TRAMA while conditioning on the true ARG.

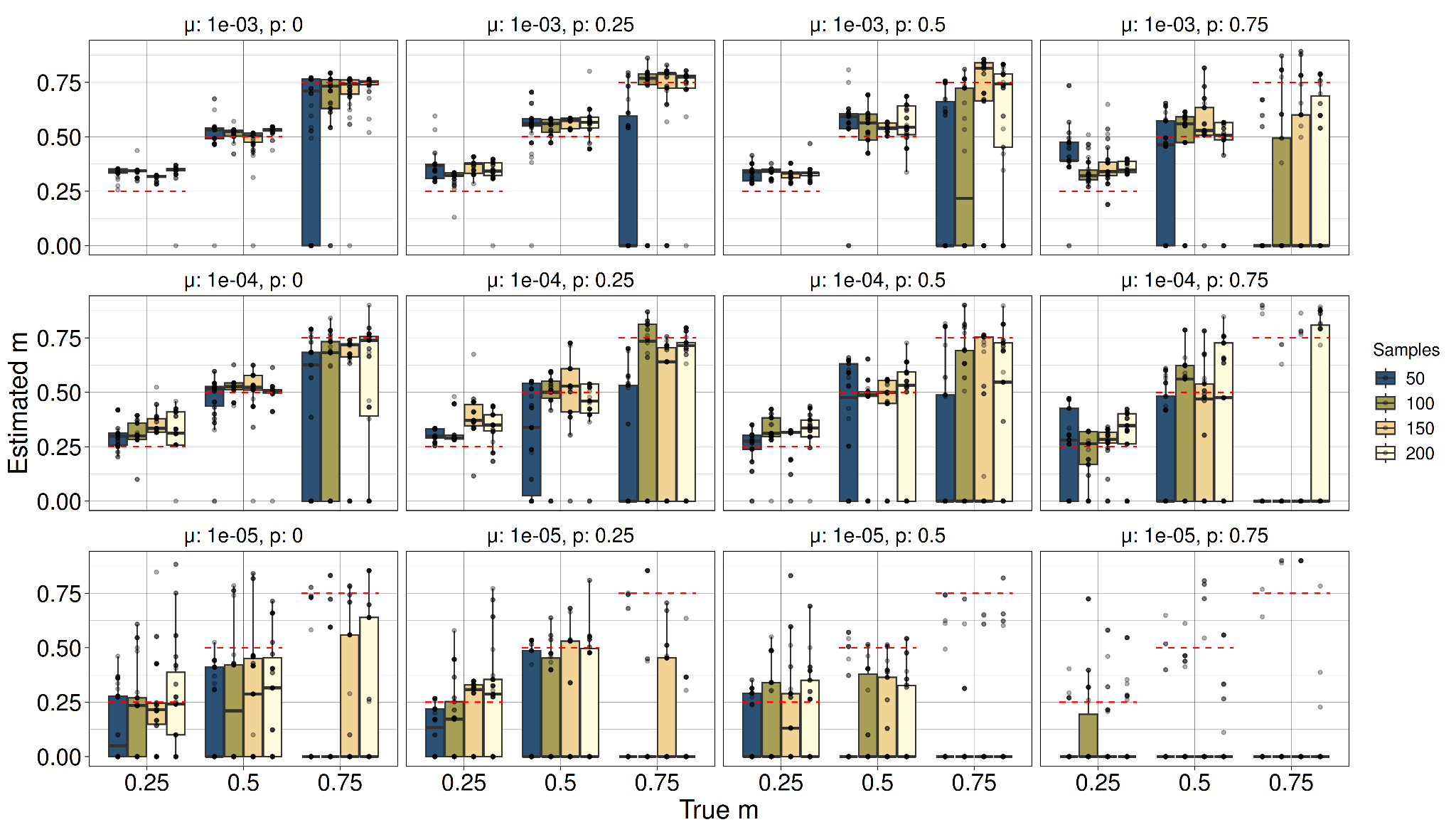

**Figure S1C. Intersectional accuracy of the multistep mutation length parameter (*m*) under the Two-Phase Mutation Model (TPM).** Accuracy of *m* estimation was evaluated across different values of μ, *p*, *m*, and sample sizes. TR variants were simulated in independent simulations under the TPM, and parameters were estimated using TRAMA while conditioning on the true ARG.

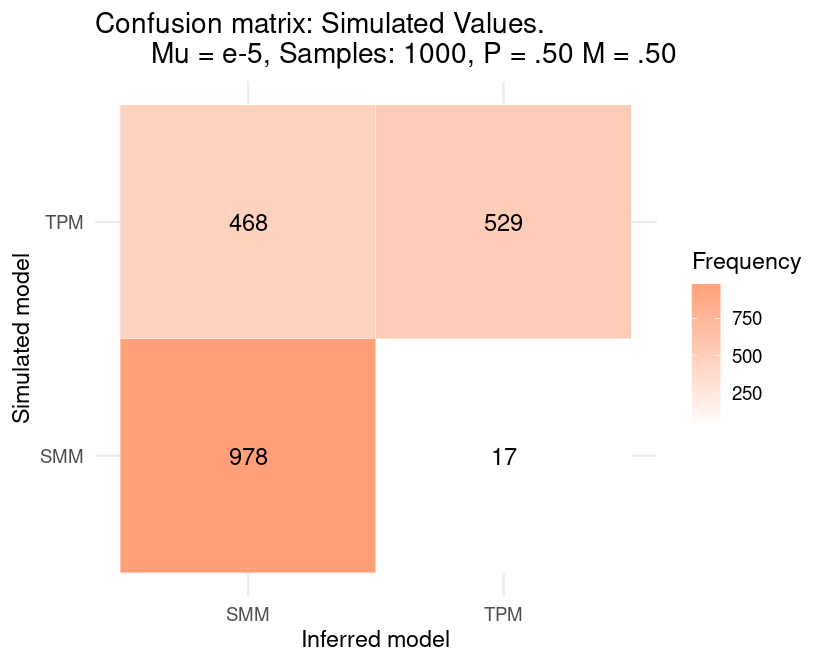

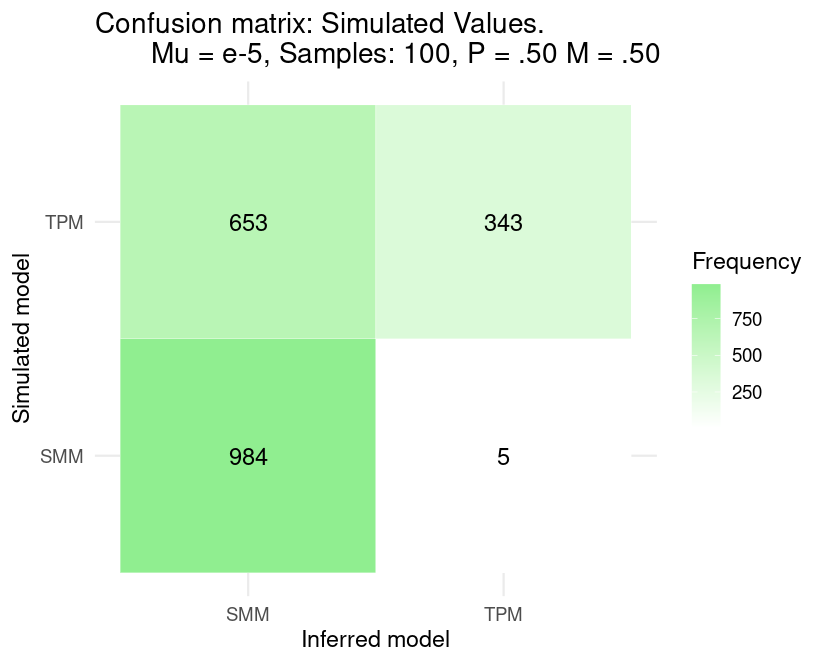

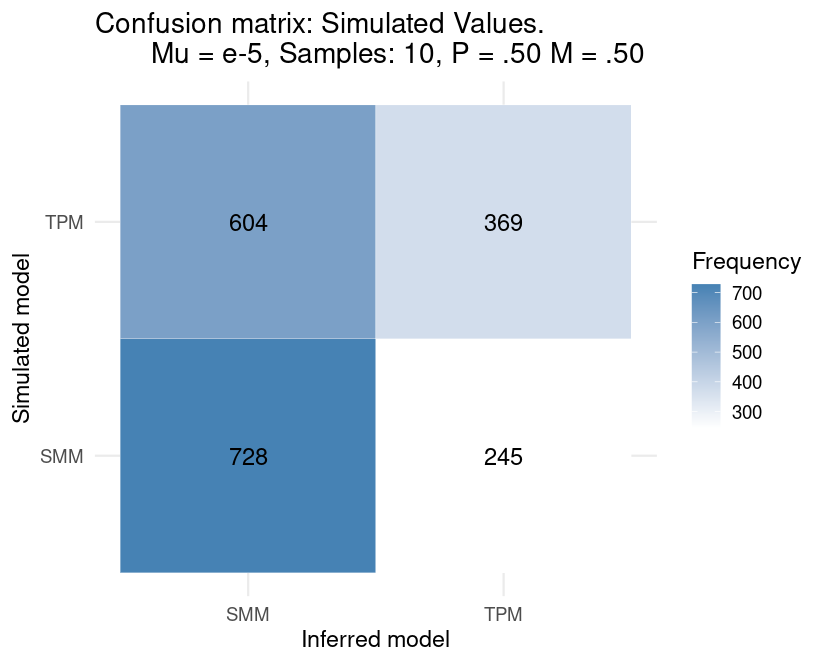

**Figure S2A**.- Confusion matrices for 3 different sample sizes (N = 10, 100 and 1000) showing how TRAMA classifies 1,000 simulations as evolving either under the SMM or TPM using a mutation rate of μ = 10^-5^.

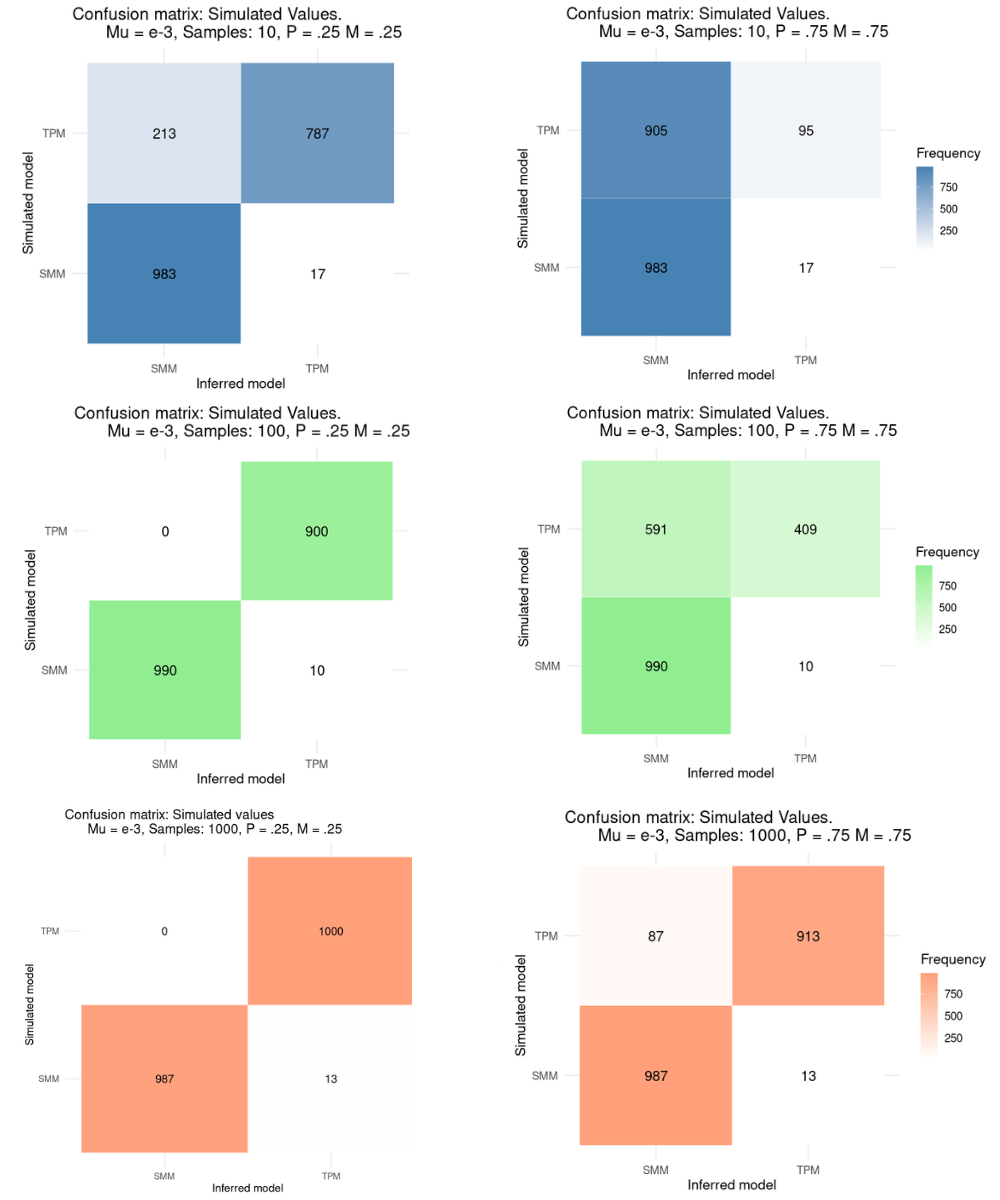

**Figure S2B**.- Confusion matrices for 3 different sample sizes (N = 10, 100 and 1000) showing how TRAMA classifies 1,000 simulations as evolving either under the SMM or TPM using a mutation rate of μ = 10^-3^, and a p = m = .25 and .75 for the TPM.

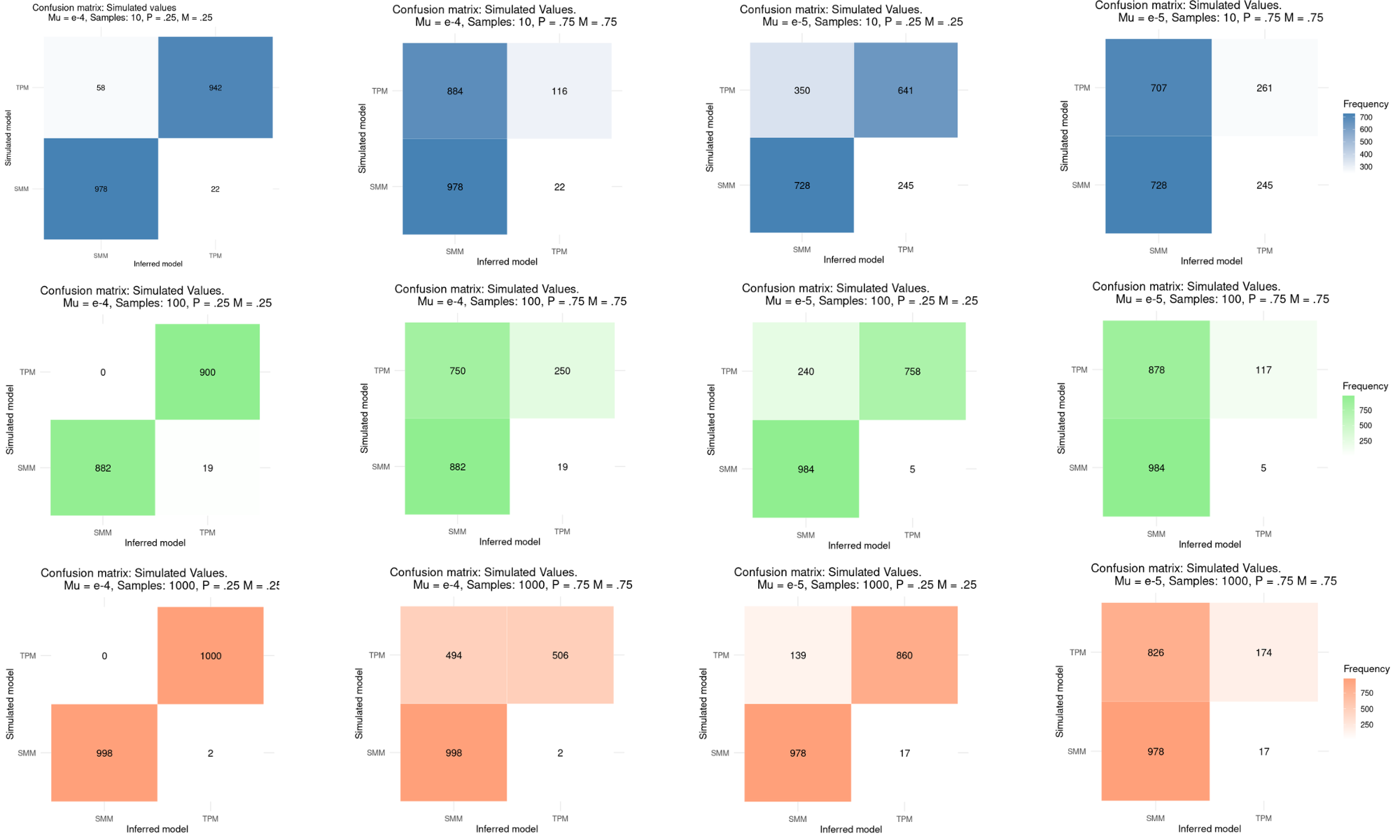

**Figure S2C**.- Confusion matrices for 3 different sample sizes (N = 10, 100 and 1000) showing how TRAMA classifies 1,000 simulations as evolving either under the SMM or TPM using a mutation rate of μ = 10^-4^  and 10^-5^ , and a p = m = .25 and .75 for the TPM.

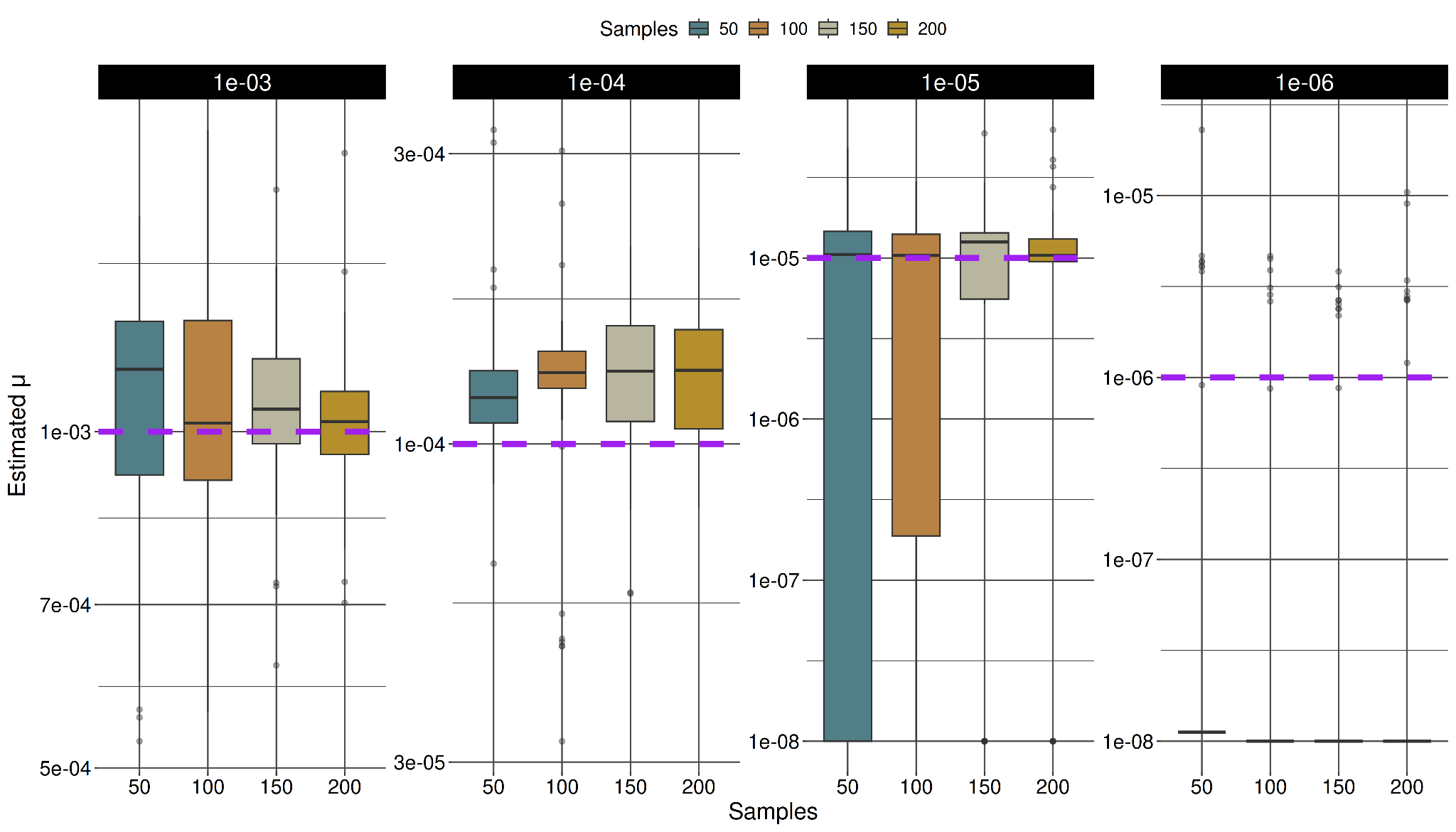

**Figure S3. Estimation of mutation rate under the Stepwise Mutation Model (SMM) using an inferred genealogy.**

Tandem repeat (TR) variants were simulated in independent simulations for each mutation rate (μ = 10⁻² to 10⁻⁶) under the SMM. For each simulated dataset, μ was estimated using TRAMA across multiple sample sizes (N = 50, 100, 150, and 200 individuals), conditioning on an ancestral recombination graph estimated from the data.

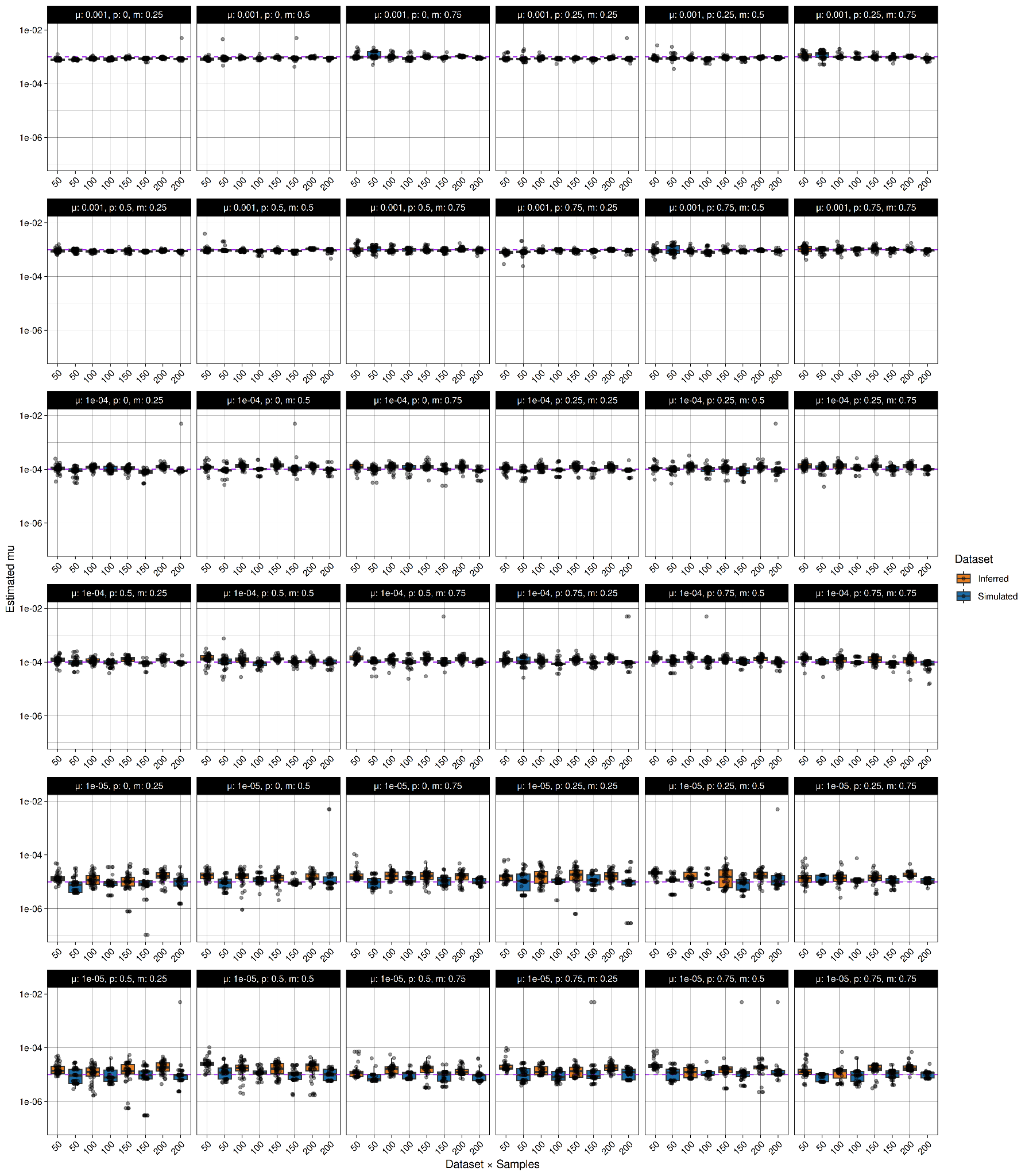

**Figure S4A. Comparison of mutation rate (μ) estimation under the Two-Phase Mutation Model (TPM) using true versus estimated genealogies.** Mutation rate (μ) estimation accuracy was evaluated across different values of μ, *p*, *m*, and sample sizes under the TPM. TR variants were simulated in independent simulations, and μ was estimated using TRAMA while conditioning on either the true genealogy or an estimated genealogy. Estimates based on estimated genealogies show increased variance across parameter values relative to those conditioned on the true genealogy.

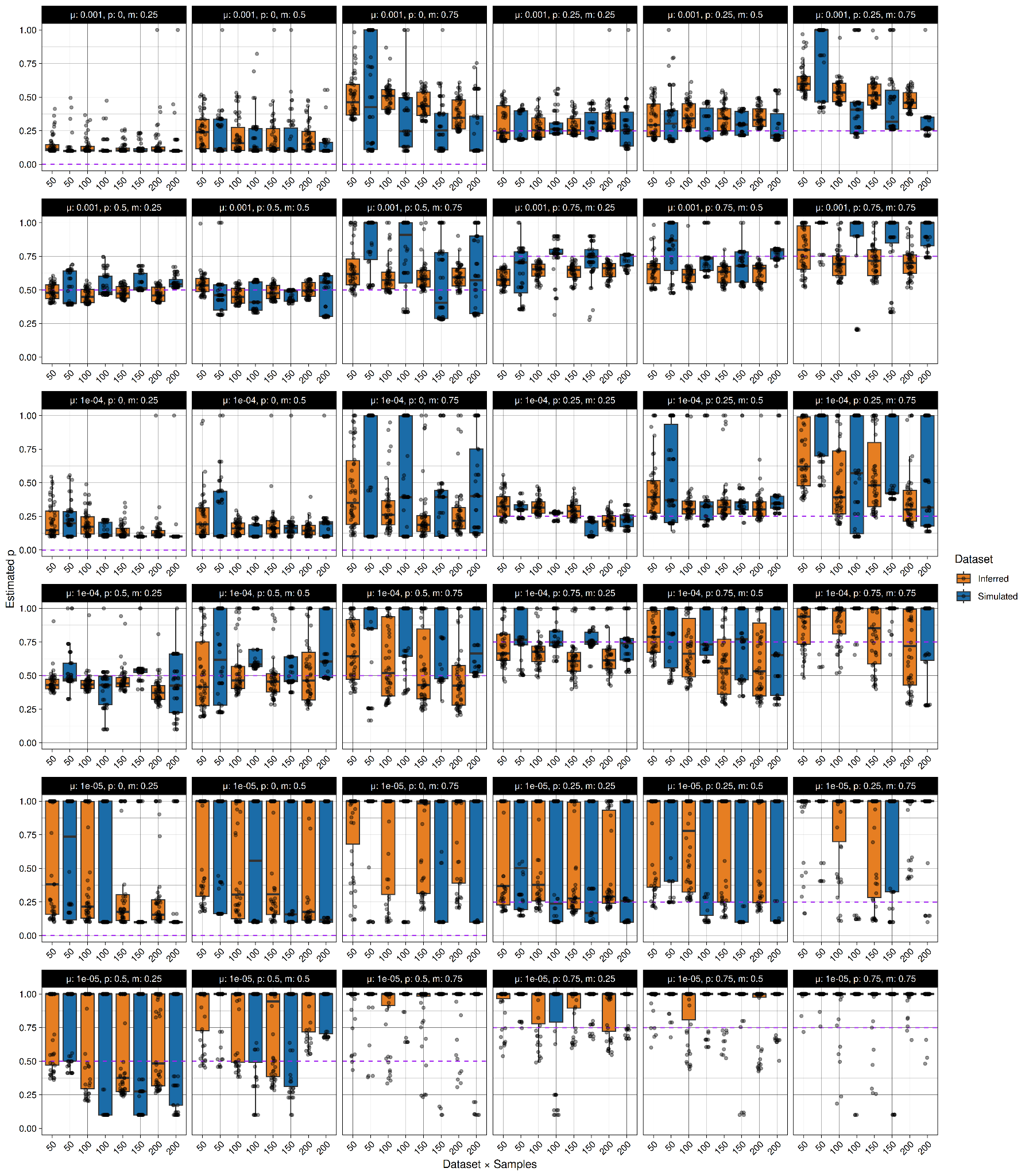

**Figure S4B. Comparison of multistep mutation probability (*p*) estimation under the Two-Phase Mutation Model (TPM) using true versus estimated genealogies.** Accuracy of *p* estimation was assessed across combinations of values of the mutation rate (μ), multistep mutation probability (*p*), multistep mutation length (*m*), and sample sizes . Parameter estimates were obtained using TRAMA while conditioning on either the true genealogy or an estimated genealogy.

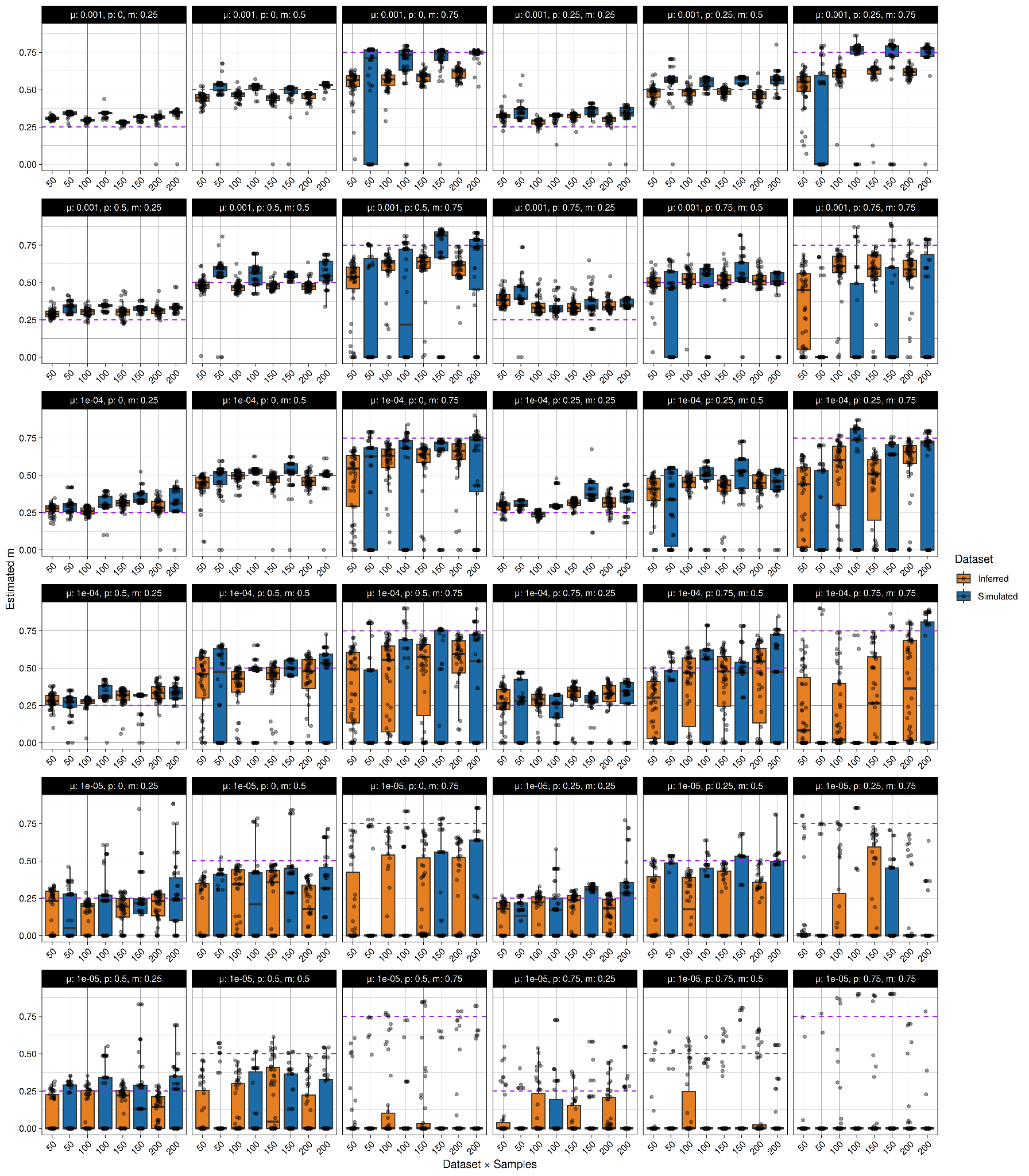

**Figure S4C. Comparison of multistep mutation length (*m*) estimation under the Two-Phase Mutation Model (TPM) using true versus inferred ARGs.** Accuracy of *m* estimation was evaluated across different values of μ, *p*, *m*, and sample size (N = 10–1,000). TR variants were simulated in independent simulations under the TPM, and parameters were estimated using TRAMA while conditioning on either the true genealogy or an estimated genealogy.
